# Topological adaptation of heliorhodopsins enables exogenous second-antenna acquisition in monoderm phototrophs

**DOI:** 10.1101/2025.11.27.691015

**Authors:** Daniil Kornilov, Sergey Bukhdruker, Roman Astashkin, Fedor Tsybrov, Mikhail Shevtsov, Siarhei Bukhalovich, Sergey Ivashchenko, Anatolii Mikhailov, Egor Zinovev, Vitaliy Golybev, Yury Ryzhykau, Alexey Vlasov, Maksim Rulev, Valentin Borshchevskiy, Pavel Kuzmichev, Ernst Bamberg, Valentin Gordeliy

**Affiliations:** Moscow Center for Advanced Studies, Kulakova str. 20, Moscow 123592, Russia; Department of Chemical Engineering and Chemistry, Laboratory of Self-Organizing Soft Matter, Eindhoven University of Technology; Eindhoven 5612 AP, the Netherlands; Institute for Complex Molecular Systems; Eindhoven University of Technology, Eindhoven 5612 AP, the Netherlands; Frank Laboratory of Neutron Physics, Joint Institute for Nuclear Research; Dubna, Russia; Department of Cell and Molecular Biology, Biomedical Centre, Uppsala University; 75124 Uppsala, Sweden; Department of Biophysical Chemistry, Max Planck Institute of Biophysics; 60438 Frankfurt am Main, Germany; Institut de Biologie Structurale Jean-Pierre Ebel, Université Grenoble Alpes–Commissariat à l’Energie Atomique et aux Energies Alternatives–CNRS; F-38027 Grenoble, France

## Abstract

Heliorhodopsins (HeRs), the third rhodopsin family, are characterized by inverted membrane topology and confinement to monoderm organisms, yet their biological meaning has so far remained a mystery. We report the first crystal structure of a eukaryotic HeR, supported by structural modeling and comparative analyses across all domains of life. A conserved carotenoid-binding site, reminiscent of secondary antennae in some microbial rhodopsins, is identified and found to be common among HeRs. We show that inverted topology allows recruitment of exogenous xanthophylls, inaccessible in diderm cells, explaining HeRs’ distinctive orientation and distribution. These findings reveal a previously unrecognized light-harvesting mechanism of HeRs, expand the known repertoire of microbial phototrophy, and suggest evolutionary constraints linking membrane topology to environmental metabolite accessibility.

## Introduction

Rhodopsins play a crucial role in life on Earth and more specific are central to optogenetics. Heliorhodopsins (HeRs) are an unusual group (Type III) of microbial rhodopsins recently discovered through functional metagenomics ^1^. Members of this group exhibit a genetic divergence from the known (Type I and II) rhodopsins, but share a common feature: seven transmembrane helices with a conserved lysine residue in the seventh helix, typical of all rhodopsins. However, their distinct feature is that they exhibit their N- and C-terminal tails inside and outside the cytoplasm, respectively, which is the opposite orientation of all previously known rhodopsins ^1,2^.

Bioinformatics studies have shown that HeRs are widely distributed across multiple domains of life, including archaea, bacteria, and eukaryotes, and can be classified into several distinct subgroups ^1,2^. Together, they constitute approximately 20% of all known microbial rhodopsins ^3^. Moreover, organisms harboring HeRs occur in diverse environmental niches ^1,4^, indicating a potentially broad range of biological functions. Interestingly, despite this wide phylogenetic and ecological distribution, HeR genes are predominantly found in organisms that lack an outer membrane ^4,5^. This pattern contrasts with the broader taxonomic presence of Type I rhodopsins, which are found across many organisms, including those with an outer membrane.

It has been suggested that the absence of HeRs in diderms may be related to the structural and functional properties of the outer membrane (OM) in diderm bacteria ^4^. The OM, an asymmetric bilayer of lipopolysaccharides and glycerophospholipids, serves as a selective barrier, impeding the penetration of glycopeptides ^6^, amphiphilic compounds ^7^, and, in some cases, light when heavily pigmented ^8^. However, why HeRs are absent in diderm cells and inverted in membranes, and whether these distinctions are related, remain central mysteries in the rhodopsin world.

We hypothesize that identifying specific structural features common to all HeRs may help to resolve these mysteries. Although a number of HeRs have been identified, only a few have been partially in detail studied. Among them there are 48C12 ^1,2,9–12^, *Thermoplasmatales archaeon* SG8-52-1 heliorhodopsin (*T*aHeR) ^10,12,13^, *Bellilinea caldifstulae* heliorhodopsin (*Bc*HeR) ^14^, *Actinobacteria bacterium* IMCC26103 heliorhodopsin (*Ab*HeR) ^15^, *Trichococcus flocculiformis* heliorhodopsin (*Tf*HeR) ^16^, and *Omithinimicrobium cerasi* heliorhodopsin (*Oc*HeR) ^17^. However, structural insights into HeRs remain scarce, with only two representative proteins having available high-resolution structures: bacterial 48C12 ^2,9,18^ (PDB IDs: 6SU3, 6SU4, 6UH3 and 9LJJ) and archaeal *T*aHeR ^13,19,20^ (PDB IDs: 7U55, 6IS6, and 7CLJ). However, the structure of a eukaryotic HeR is still missing.

To complete the structural characterization of HeRs, we solved the high-resolution structure of a HeR from *Chromera velia* CCMP2878 (UniProt ID: A0A0G4GC06) (hereafter referred to as Cv06), a representative of eukaryotic HeRs, and also performed its biophysical characterization.

Further structural analysis of the archaeal, bacterial, and eukaryotic rhodopsins revealed the presence of so-called lateral fenestration (an opening in the rhodopsin exposing the retinal β-ionone ring to the external environment) in all three proteins. It is known that fenestration is unique to Type I rhodopsins, which have a second chromophore (carotenoid) antenna on the protein’s surface. Examples include xanthorhodopsin (XR) from the extreme halophilic bacterium *Salinibacter rubeus* with salinixanthin as the second antenna ^21,22^, rhodopsin from *Gloeobacter violaceus* (GR) with echinenone ^23,24^, rhodopsin Kin4B8, which can form complexes with lutein and zeaxanthin ^25^, and heimdallarchaeaial rhodopsins ^26^. The discovery of numerous genes of microbial rhodopsins across various domains of life highlights a general structural requirement: the specific fenestration in the rhodopsin molecule that facilitates carotenoid antenna binding.

Motivated by these findings, we conducted bioinformatic and structural AlphaFold 3 analysis of all available HeRs data and found that nearly all of them comprise fenestrations. These results demonstrate that HeRs are the two-antenna rhodopsins. However, the question remains as to why they are exclusively present in monoderm cells. In our work, we show that this is explained by the fact that the outer membrane of Gram-negative bacteria is impermeable to carotenoids, which serve as the second antenna. This observation supports the hypothesis that HeRs represent a type of rhodopsins capable of acquiring xanthophylls from the extracellular environment. Assuming that HeRs depend on carotenoids sourced from the environment, their inverted orientation within the membrane relative to other rhodopsins can also be explained. This is due to the geometrical constraints imposed by the size of the amphiphilic carotenoids and the asymmetrical position of the β-ionone ring of retinal relative to the geometric center of the lipid bilayer (and the protein).

Further, we present our data, their analysis, and emphasize, that HeRs provide a significant advantage for phototrophy on Earth.

## Results

### Expression and characterization of eukaryotic heliorhodopsin Cv06 from Chromera velia

To determine common structural features of HeRs, which could help to disclose the central mysteries of their nature, we first decided to complete the available structures of archaeal and bacterial HeRs with those of eukaryotic ones.

Not all microbial rhodopsins can be expressed well in standard expression systems like *E. coli*. To achieve efficient expression of the eukaryotic Cv06, we employed the LEXSY system, following previously established protocols for channelrhodopsin 2 from *Chlamydomonas reinhardtii* alga ^27^ and a proton-pumping rhodopsin from *Leptosphaeria maculans* fungi ^28^. Expression of Cv06 was described in detail in our work ^29^.

pH titration experiments were conducted to determine the pK_a_ values of the retinal Schiff base and its counterion (Fig. S1). However, due to the instability of the protein at highly acidic and basic pH levels, precise pK_a_ values could not be determined. Fig. S1 illustrates that the pKa values for the counterion and the Schiff base are likely below 4.5 and above 11, respectively. This suggests that the Schiff base remains protonated over a wide pH range. This result is very close to that for 48C12 ^1,2^, *Bc*HeR ^14^, and *T*aHeR ^13^.

### Black lipid membrane assay

Certain microbial rhodopsins exhibit a light-dependent ion transport capability. However, in the case of many characterized HeRs, hydrophobic residues occupy the extracellular half of these proteins ^2^, blocking the entry pathways for ions. To verify whether this is also the case for Cv06, we conducted black lipid membrane (BLM) assays on Cv06 within lipid vesicles at pH 7.2 (Fig. S2). We used Bacteriorhodopsin from *Halobacterium salinarum* (BR) as a positive control.

Beyond small ions, which are typically transported by known rhodopsins (e.g., H^+^, Na^+^, K^+^, and Cl^-^), we also explored the Cv06 potential transport of larger ions, such as acetate, bicarbonate, and nitrate ions. This was motivated by the presence of such molecules in the structures of 48C12, located within the hydrophilic cavity near the Schiff base at acidic pH ^2^ and neutral pH ^18^, which can move between the hydrophilic cavity and cytoplasm ^30^. Our experiments revealed no ion transport activity for Cv06. The absence of transport activity has been also observed for 48C12 ^1,2^ and *T*aHeR^13^, while proton transport activity has been shown for viral HeRs ^31^.

### Photocycle

We performed time-resolved flash-photolysis experiments with Cv06 at pH 7.5 in a temperature range of 20 – 40°C. Exposure of microbial rhodopsins to a brief flash of light triggers the isomerization of the retinal molecule. This event initiates a sequence of reactions known as the photocycle ^32,33^. In the case of HeRs, it extends over several seconds ^1,13,14^. Typically, a photocycle includes various intermediates, denoted by Latin letters, with their properties analyzed by analogy with bacteriorhodopsin ^33^. Fig. 1a presents the time-dependent absorbance difference of Cv06, covering a wavelength range from 330 nm to 730 nm at a temperature of 20°C. The photocycle becomes faster with increasing temperature (Fig. 1b). A global multiexponential fit and a model of irreversible transitions revealed temperature dependence of decay half-times, thermodynamic parameters of reactions (Fig. 1c), and allowed the obtaining of the photocycle model (Fig. 1d), intermediate spectra (Fig. 1e).

**Figure 1.**
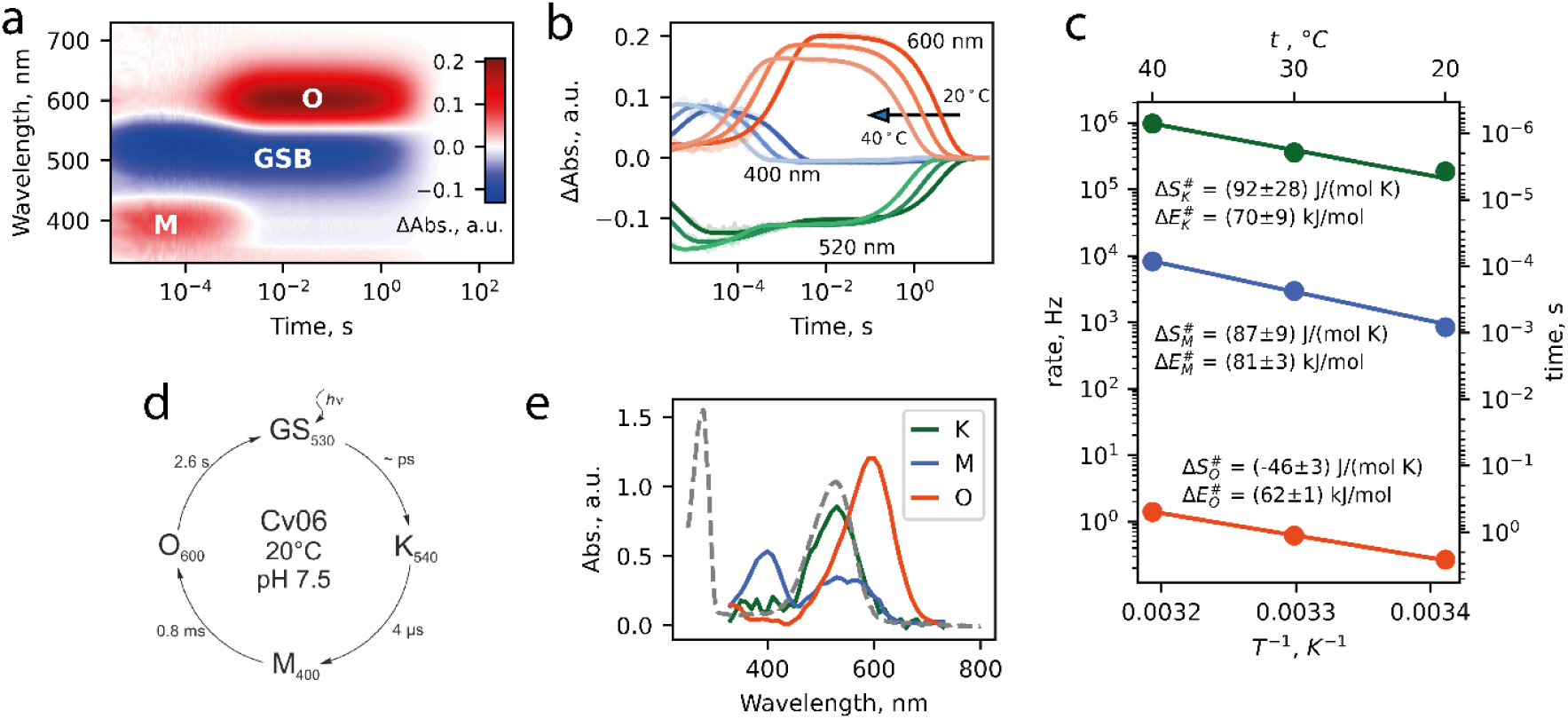
Time-resolved flash-photolysis measurements. Cv06 protein solution was suspended in the buffer 20 mM Na-phosphate (pH 7.5), 500 mM NaCl, 10% glycerol, 0.05% (w/v) DDM. (**a**) Transient absorption changes of Cv06 in DDM micelles excited at 520 nm at 20°C. Here, M and O denote the areas corresponding to the M and O states, GSB states for ground state bleaching. (**b**) Time evolutions of transient absorption changes at specific wavelengths of Cv06 in DDM micelles and their temperature dependence (bright lines - fitted data, faded - measured data). (**c**) Half-times temperature dependence shown in Arrhenius coordinates with estimated enthalpy (ΔE#) and entropy (ΔS#) of the transitions. (**d**) The model of the Cv06 photocycle. Indices denote the absorption maximum of intermediate states. (**e**) Reconstructed absorption spectra of the intermediates obtained by the multi-exponential global fitting and the model of irreversible transitions. The dashed line shows the absorption spectra of the ground state of Cv06.

Reconstructed photocycle, absorption spectra of intermediates, and half-times of intermediate states are close to those of 48C12 ^2^ and *T*aHeR ^13^. After a flash laser pulse, K state, the first revealed state, decays with a half-time of 4 us into the blue-shifted M state. The M state with absorption maximum of 400 nm corresponds to the retinal Schiff base deprotonation in the photocycle of microbial rhodopsins. This blue-shifted M state decays into a long-lived, red-shifted O state with a half-time of 0.8 ms. Such a long-lived O state is a distinctive feature of bacterial (48C12) and archaeal (*T*aHeR) HeRs. Our data for eukaryotic Cv06 shows that this feature is spread among HeRs from all domains of life. The half-time of several seconds of the O state decay to the ground state could indicate the presumable sensory function of this HeRs ^1^. Notably, *T*aHeR exhibits a slightly prolonged decay time to the ground state, as discussed in ref. ^34^. Arrhenius plot for Cv06 shows a linear dependence of half-times, and the value of enthalpy of transitions is quite close to the other microbial rhodopsins, e.g., bacteriorhodopsin ^35^.

### Сrystallization

Crystallization of Cv06 was thoroughly described in ref. ^29^. Briefly, the protein was crystallized *in meso* similarly to our previous works ^36–38^. The initial crystals were pseudo-merohedral twins, which prevented structural solution. Twinning was mitigated by utilizing monopalmitolein instead of monoolein as an LCP lipid and lowering the concentration of ammonium phosphate buffer used as precipitant ^29^. The resultant crystals contained a low twinning fraction, yet their low diffraction resolution (with anisotropic cutoffs of 3.2×3.0×4.1 Å) prevented further structural refinement.

In this work, we further improved crystal quality by changing the precipitant from 2.0 M ammonium phosphate buffer (pH 7.6) to 1.2 M sodium potassium phosphate buffer (pH 8.2). This exchange further slowed the crystal growth (to about 4 months), which positively affected the diffraction resolution. As a result, the resolution was improved to 2.5×2.4×3.0 Å, which allowed us to build an atomic model of Cv06.

### Structure overview

In the crystal structure, one asymmetric unit (ASU) contains four Cv06 dimers. Eight monomers, denoted as chains A–H, are almost identical in structure (with root mean square deviation, rmsd, between Cα atoms not exceeding 0.085 Å). The structure of Cv06 in comparison with other known HeR structures (48C12 ^2^ and *T*aHeR ^13^) shows that the overall fold is conserved with minor peculiarities. The significant differences are shown in Fig. S3. First, helices A and D are 7 Å longer in Cv06 in the cytoplasmic side. Second, the short helix between helices B and C in 48C12 deflects from that in *T*aHeR and Cv06 with a rmsd of 0.46 Å. Third, AB-loop (loop between helices A and B) in Cv06 has about 14 Å longer β-sheets in comparison with 48C12 and *T*aHeR, and is bordered by one coil of α-helix that is absent in 48C12 and *T*aHeR. Finally, helices E in all HeRs deflect from each other in the helix’s outer half with a maximal rmsd of 0.44 Å.

One of the key features of HeRs, dimerization, is also preserved in Cv06. However, the dimerization interface exhibits several differences from that of 48С12 (Fig. S4). Firstly, the extracellular part of the dimer is stabilized not by a single hydrogen bond (Y179-D127’ in 48С12) but by two - Y229’ of helix E simultaneously connects to E171 and S173 of helix D, which replace D127’ and V129’, respectively. Additionally, the extracellular part contains only one hydrophobic pair (V225’-V176) instead of two (I178-V129’ and I174-L132’) found in 48С12. Secondly, the cytoplasmic half of the protein lacks stabilizing hydrogen bonds that support dimerization. In 48С12, Y151 of helix D forms a bond with D158’ of the DE-loop, whereas in the case of Cv06, Y151 is replaced by H195, which forms a hydrogen bond with D208 (corresponding to D158 in 48C12) within the same protomer. However, the intracellular part of the protein is further stabilized by two hydrophobic pairs A198-L199’ of helices D, resulting from the extended helix D.

As was previously suggested, the topology of membrane proteins follows the “positive inside” rule ^39^. In contrast, HeRs follow the “positive inside and negative outside” rule rather than just “positive inside” ^2^. In 48C12 and *T*aHeR this rule is well expressed. In Cv06 (Fig. S5), both inner and outer sides of the protein have almost an equal number of positively and negatively charged residues; nevertheless, the predominance of negatively charged residues by one in the outer side and positively charged residues by 2 in the inner side show that the rule is also valid for Cv06.

### Analysis of the hydrogen-bonded networks in HeRs

The extracellular half of the protein consists exclusively of hydrophobic residues, with the exception of T115. T115 has no opportunity to form large chains of hydrogen bonds with its surrounding ^36,37,40^ (its only hydrogen is donated to the carbonyl group of F111), indicating that the protein is not an ion transporter. This conclusion agrees with our BLM results.

In contrast to the extracellular half of Cv06, there is a broad hydrogen bond network from the Schiff base to the cytoplasm (Fig. 2), forming three key regions – retinal Schiff base (RSB), hydrophilic cavity in the vicinity of RSB – a feature, reported for other HeRs, and a region of highly conserved E193 (E149 in 48C12 and E150 in *T*aHeR).

**Figure 2.**
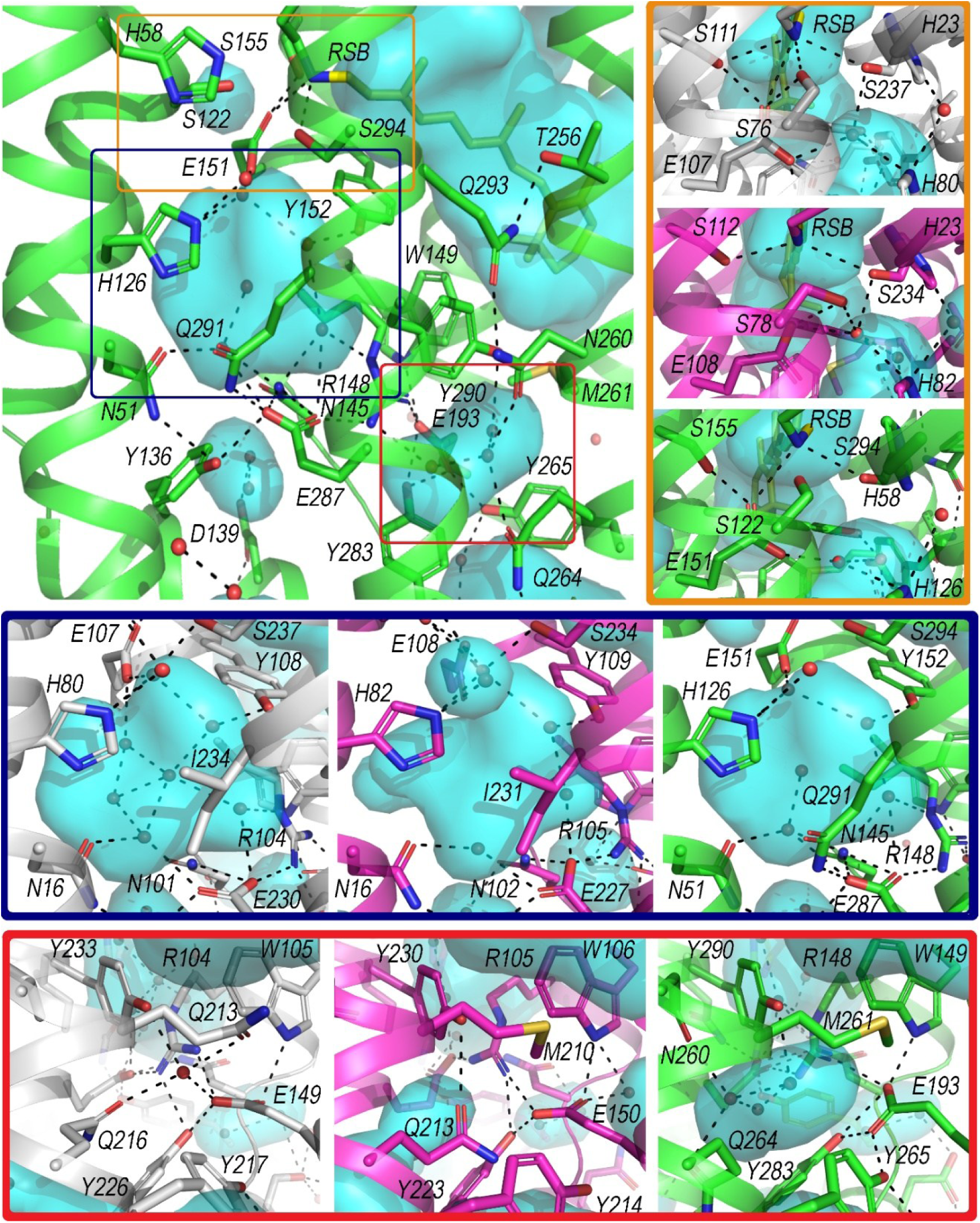
Map of intramolecular hydrogen bonds. Proton wires of Cv06 (PDB ID: 9VFC, chain A, marked green), 48C12 (PDB ID: 6SU3 ^2^, chain A, marked grey), and TaHeR (PDB ID: 6IS6 ^13^, chain A, marked magenta). Retinal cofactor marked yellow; cavities marked cyan; water molecules are shown as red spheres; hydrogen bonds marked dashed lines. Waters, hydrogen bonds, and amino acid residues in all the structures are shown for an “A” alternative conformation only.

The RSB region consists of the all-trans retinal and the hydrophilic surrounding, similar to other HeRs. RSB forming the contacts with E151 and S294 (Fig. 2, orange) in all eight protomers. It additionally forms a contact with S122 in protomer B, but does not form a bond with S155 in any of the protomers. Residues making the network around the RSB are identical in composition within these three HeRs.

One of the unique features of HeR is the presence of a large hydrophilic cavity close to the retinal Schiff base. Residues forming this cavity are highly conserved among all the HeRs, but some HeRs have distinct features. Fig. S6 shows that F103 in 48C12 and F147 in Cv06 are replaced by Y104 in *T*aHeR. Moreover, the Y104 ring in *T*aHeR is rotated approximately 55° and 65° relative to the F103 in 48C12 and F147 in Cv06, respectively. This orientation enables the formation of an additional small cavity in *T*aHeR between N102 and Y104. Next, V83 in 48C12 and V129 in Cv06 are replaced by I85 in *T*aHeR. A81 in *T*aHeR and A125 in Cv06 are replaced by F79 in 48C12. In the hydrophilic cavity, the network is the same within three HeRs with one exception (Fig. 2, dark blue). In Cv06, Q291 replaces I234 in 48C12 and I231 in *T*aHeR, forming an additional hydrogen bond with conserved E287 (E230 in 48C12, E227 in *T*aHeR), but not with waters in the cavity.

Finally, all three HeRs comprise a hydrophilic region around the conserved E193 (in Cv06) residue, albeit having some unique features. In 48C12, this region contains hydrophilic Q213, instead of hydrophobic M261 in Cv06 and M261 in *T*aHeR (Fig. 2, red). In Cv06, there is an additional cavity containing three water molecules, which are absent in the *T*aHeR structure. In contrast, the 48C12 structure contains only one water molecule in this region. If HeRs can act as enzymes as it is assumed ^2^, then the cavity near E193 could be the active site rather than the cavity near the Schiff base, since the former is more easily accessible from the cytoplasm.

The retinal environment was further examined in HeRs. Differences in the residue composition forming the retinal pocket between the three HeRs are depicted in Fig. S7. A unique feature of Cv06 in this area is the presence of two additional polar residues: S297 and Y250 instead of hydrophobic A240 and L202 in 48C12, and A237 and I199 in *T*aHeR. The presence of hydrophilic residues may be responsible for the 10 nm blue shift in Cv06’s absorption compared to 48C12 and *T*aHeR.

Accordingly, Cv06 exhibits both conserved and unique structural characteristics with its archaeal and bacterial relatives in Type III rhodopsins. However, we will demonstrate that, despite different domains of life, HeRs possess a unique common structural motif on the protein surface, known in Type I rhodopsins as *fenestration*, which facilitates the binding to their surface of xanthophylls that serve as a second antenna.

### Lateral fenestration on the surface of HeRs

One of the most noticeable features we detected in all the considered HeRs is the fenestration on its surface that allows the retinal pocket to be accessible from the outside (Fig. 3). This fenestration is similar to those found in Type I rhodopsins, such as XR ^22^, GR ^23,24^, Kin4B8 ^25^, and the sodiumium pump (NaR) ^41^, having 90% identity with KR2. In all these cases (NaR only with the single mutation T216G), structural studies have shown that similar fenestrations accommodate oxygenated carotenoids (xanthophylls), which serve as second antennas. For example, the keto-ring of salinixanthin in XR is positioned on the surface of the fenestration, indicating that it acts as a carotenoid binding site (CBS).

**Figure 3.**
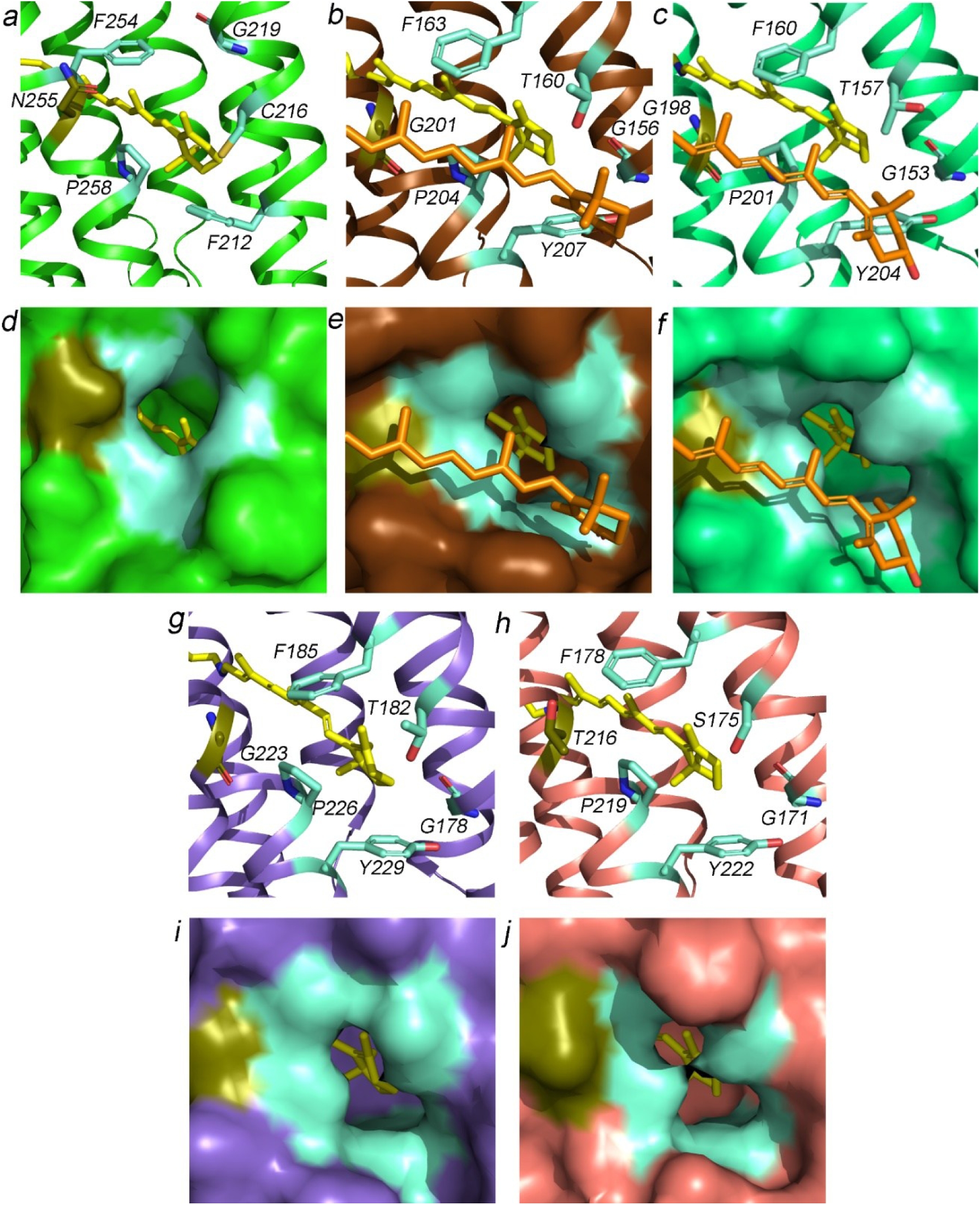
Lateral fenestration in microbial rhodopsins. Cv06 (PDB ID: 9VFC) marked green (**a**, **d**), XR (PDB ID: 3DDL ^74^) marked brown (**b**, **e**), Kin4B8 (PDB ID: 8I2Z ^25^) marked spring green (**c**, **f**), GR (PDB ID: 6NWD ^75^) marked purple (**g**, **i**) and KR2 (PDB ID: 4XTL ^76^) marked pink (**h**, **j**). Panels (**a, b, c, g, h**) show amino acid residues forming lateral fenestration, while panels (**d, e, f, I, j**) show protein surfaces. Retinal marked yellow; external carotenoids: salinixanthin for XR, panels (**b, e**), and zeaxanthin for Kin4B8, panels (**c, f**) are marked orange; amino acid residues forming the fenestration marked cyan; conserved asparagine residue in HeRs with corresponding glycine in XR and GR and threonine in KR2 marked splitpea.

In Cv06, the CBS is primarily formed by five amino acid residues: F212, C216, G219, F254, and P258 (Fig. 3 and 4). The aromatic rings of F212 and F254 are located on opposite sides of the fenestration, as well as the residues G219 and C216. Our analysis shows that both 48C12 and *T*aHeR have similar fenestrations (Fig. 4). Indeed, the differences between all the HeRs are minor: in 48C12 and *T*aHeR, P258 is replaced by A210 and A207, respectively (Fig. 4). In *T*aHeR, F212 is further replaced by Y164. When comparing this site in HeRs to the fenestration found in Type I rhodopsins with characterized carotenoid binding, the only notable difference we found is the presence of an asparagine residue (N255 in Cv06, N207 in 48C12, and N204 in TaHeR) in the position of glycine (G201 in XR, G198 in Kin4B8, and G223 in GR), where carotenoid is located (Fig. 3).

**Figure 4.**
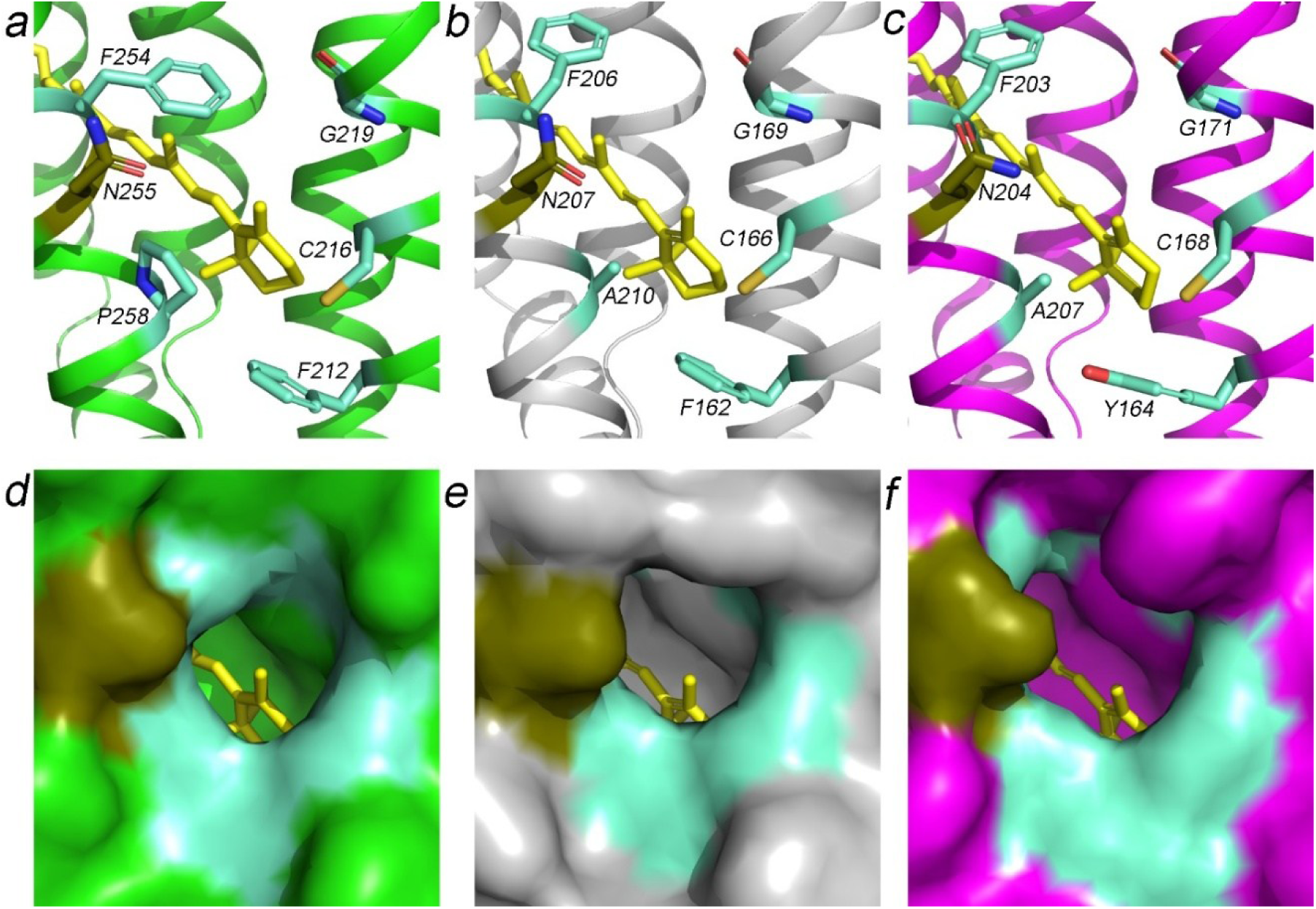
Lateral fenestration in HeRs. (**a**, **d**) Cv06 (PDB ID: 9VFC, marked green), (**b**, **e**) 48C12 (PDB ID: 6SU3 ^2^, marked gray), and (**c**, **f**) *T*aHeR (PDB ID: 6IS6 ^13^, marked magenta). Panels (**a-c**) show amino acid residues forming lateral fenestration, while panels (**d-f**) show protein surfaces. Retinal marked yellow; amino acid residues comprising the fenestration marked cyan; conserved asparagine residue in HeRs marked splitpea.

Thus, although the eukaryotic Cv06 shares both common and unique structural features with its archaeal and bacterial counterparts among HeRs, all possess a fenestration. This fenestration enables the binding of xanthophylls to their surface, which function as secondary antennas. The organization of this site is highly similar to that found in Type I rhodopsins, for which the binding of the second antenna was confirmed.

### Heliorhodopsins are two-antenna proteins

Three HeRs with known structures show the presence of fenestration, and they come from different domains of life. While this did not imply that all HeRs are two-antenna proteins, it motivated us to explore whether the fenestration is present in other HeRs.

To assess the prevalence of carotenoid-binding sites among HeRs in comparison with Type I rhodopsins, we analyzed their primary sequences from the InterPro ^42^ database, specifically searching for CBS motifs similar to that of XR, allowing for variations that permit the fenestration to exist. Analysis of primary sequences and AlphaFold3 modeling revealed that, with only limited exceptions, the occurrence of opposing aromatic amino acid residues in positions 162 and 206 in HeRs and 163 and 207 in Type I rhodopsins (Fig. 5) – is a key factor for the formation of fenestration. At the same time, the absence of bulky amino acids at positions 166 and 210 in HeRs and 160 and 204 in Type I appears to be sufficient. In HeRs, the presence of glycine at position 169 shows a strong correlation (98.5% of all sequences) with the simultaneous occurrence of the opposing aromatic pair, whereas in Type I rhodopsins this does not seem to be strictly required, as previously shown ^25^. However, in Type I rhodopsins, we observed another strict correlation at position 204, where 99.1% of all sequences with fenestration contain a proline residue.

**Figure 5.**
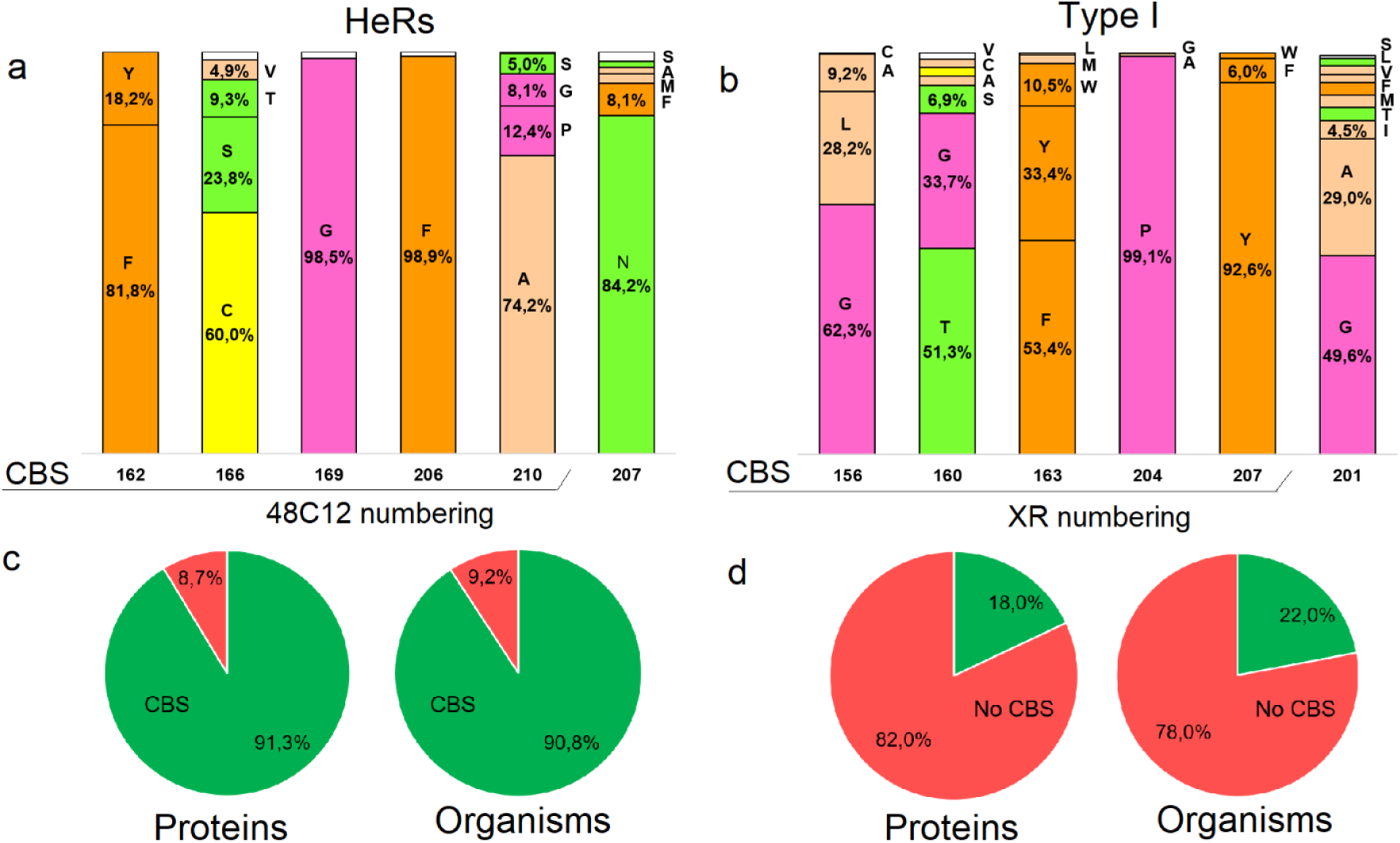
The occurrence of CBS among HeRs and Type I rhodopsins. (**a**) Composition of CBS in HeRs. (**b**) Composition of CBS in Type I rhodopsins. In (**a**) and (**b**), amino acid residues are depicted in the Zappo coloring scheme. Presence of CBS among full-size sequences of (**c**) HeRs (PF18761 InterPro ^42^ family), (**d**) Type I rhodopsins (PF01036 InterPro ^42^ family), and organisms harboring them.

Our findings show that while 18% of Type I rhodopsins possess a CBS, this feature is present in 91.3% of HeRs (Fig. 5). This significant difference suggests that HeRs function as carotenoid-binding rhodopsins (CBRs), the “two antenna rhodopsins”. Notably, subfamily 2 of HeRs is composed predominantly of viral HeRs ^2^—falls entirely within the HeR group lacking CBS domains (5% of all HeRs). Ion transport activity has been demonstrated for several members of this subfamily ^31^. The molecular nature and functional diversity of these CBS-lacking HeRs remain intriguing and warrant further investigation. One of possible explanations is that to provide inward ion transport the nature, for instance, can use topologically inverted an outward proton transporting rhodopsin. The retinal Schiff base in the case of the ground states of rhodopsin ion pumps usually should look in the direction of ion transport ^36,40,43^.

Binding of carotenoids to rhodopsin requires more than just a fenestration near the ionone ring of retinal; a suitable site on the protein surface is also necessary to accommodate the hydrophobic tail of the carotenoid. In XR, G201 allows salinixanthin tail layout on the XR surface. In addition to XR, the sodium transporter (NaR) of *Dokdonia eikasta*, which has 90% identity with KR2, has the same lateral fenestration ^41^ (in the Fig. 3 we provide the structure of KR2 due to its availability and high identity with NaR). In it, at a position corresponding to G201 in XR, there is a threonine residue, which, when replaced with glycine, allowed NaR to bind the carotenoids echinenone and canthaxanthin with the possibility of transferring energy from them to retinal ^41^. In case of HeRs and Cv06, particularly at the corresponding position, there is an asparagine residue. To assess whether, in this instance as well carotenoids can be positioned near the CBS such that their rings are proximate to the retinal ring and their hydrophobic tails align with high affinity on the protein surface, we conducted molecular docking studies with AlphaFold 3 model of Cv06 and carotenoids which could be found in environment of *Chromera velia* ^44–47^. Fig. S8 shows the most relevant variants of carotenoid layout on the surface of Cv06. For a reference comparison, we performed docking analysis of the Kin4B8 protein with the zeaxanthin molecule, whose binding has been structurally and functionally validated. Moreover, zeaxanthin shares greater structural similarity with the carotenoid molecules selected for Cv06 than salinixanthin in XR. For binding poses closely resembling that of zeaxanthin in the Kin4B8 structure (PDB ID: 8I2Z), the binding energies fall within the reference range of –11 to –10 kcal/mol, which we used as a benchmark. In our analysis, poses analogous to the zeaxanthin binding site in Kin4B8 showed binding energies in the range of –9.9 to –9.5 kcal/mol, except the peridinin molecule, which had a slightly higher value of –8 kcal/mol. Zeaxanthin exhibited the lowest binding energy (–9.9 kcal/mol), placing it very close to the reference binding range. The predicted positions we determined for selected carotenoids with Cv06 are also very similar to the position of myxol relative to NM-R3 in the Cryo-EM structure ^48^.

The absence of precise matching with the reference values is expected. It may be attributed to the lack of a pre-formed binding groove on the protein surface – a structural feature known to arise upon rhodopsin–carotenoid interaction, but which is absent in the apo form of the protein. Another evidence supporting the localization of carotenoids on the surface of Cv06 at the fenestration is the lipid molecules in our resolved protein structure. These molecules adhere tightly to the surface, including the fenestration region, and are observed in seven out of the eight protomers within the crystal lattice.

To assess whether carotenoids can consistently bind with high affinity at the fenestration site of any rhodopsin, we performed reference docking studies of bacteriorhodopsin (which lacks CBS) with zeaxanthin and canthaxanthin. Our results showed no binding poses resembling the reference carotenoid Kin4B8. The binding energies of the poses closest to the reference did not exceed –7.4 kcal/mol. Among all generated poses, the most energetically favorable were located far from the site of interest and exhibited binding energies no stronger than –8.6 kcal/mol, which are substantially weaker than the reference values.

As shown in Fig. S8, the carotenoids pass between the N255 and I259 residues. Moreover, N255 (treated as flexible) underwent almost no conformational changes during docking. These findings suggest that the presence of glycine instead of N255 is not strictly necessary for carotenoid binding, and that N255 may instead be important for properly positioning the carotenoid within the CBS.

Thus, fenestration is a common feature for the HeRs family. This is a key to resolving both mysteries of these proteins.

### The exclusive presence of HeRs in monoderm cells can be rationalized by the exogenous nature of their second antenna

Two membranes surround diderm cells. A distinct property of their outer membrane is that it is impermeable to even relatively small molecules, such as amphiphilic drugs with a molecular cut-off of around 600 Da ^49–51^. The outer membrane of Gram-negative bacteria is impermeable to carotenoids. In *E. coli*, a Gram-negative bacterium, exogenous xanthophylls cannot be acquired and therefore cannot be used as the second antenna by the expressed Type I microbial rhodopsins. Spheroids or specific *E. coli* strains with a leaky outer membrane are used to overcome the problem ^52^. It is also well known that there is a general molecular weight (MW) cut-off around 600 Da for the permeability of small hydrophilic molecules through the outer membrane (OM) of diderm cells ^53^. This corresponds to the MW of xanthophylls ^47^.

Therefore, the answer to this mystery seems straightforward if we assume that HeRs are the rhodopsins that use exogenous xanthophylls as the second antenna. This also implies that Type I CBRs in diderm cells use xanthophylls synthesized intracellularly. In contrast to Gram-negative (diderm) cells, monoderm cells in general and, in particular, Gram-positive (monoderm) bacteria may have a significant advantage due to their ability to uptake carotenoids from the environment.

### Why are the N- and C-termini inverted in membranes compared to all other rhodopsins?

While the considerations above show that HeRs are rhodopsins with carotenoids as a second antenna and explain why they are exclusively found in monoderm cells, they do not resolve the second mystery: the fact that HeRs are inverted in the membranes. This mystery should also be explained within the same conceptions, if the latter pretends to be consistent. In our case, if it is accepted that monoderm HeRs acquire their second antenna from the environment, then they should be inverted in the membranes. This is due to the geometrical constraints imposed by the size of the amphiphilic carotenoids and the asymmetrical position of the β-ionone ring of retinal relative to the geometric centre of the lipid bilayer (and the protein) (Fig. 6). Moreover, the same conception is consistent with the fact that in diderm cells, where only Type I rhodopsins are found, carotenoids are synthesized in the cytoplasm and, in contrast to environmental carotenoids, are delivered to the membrane from inside the cells ^54^.

**Figure 6.**
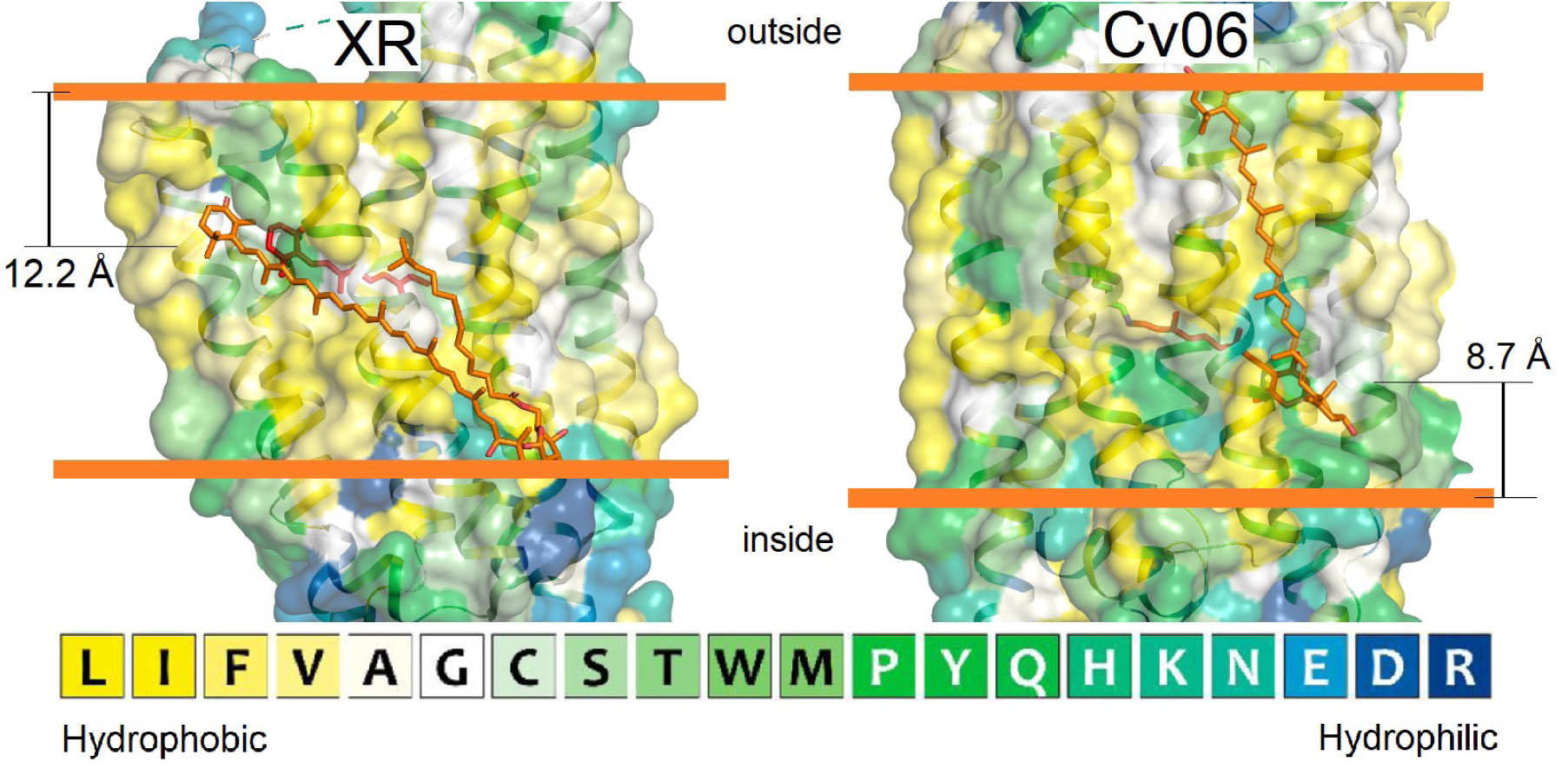
Prediction of the hydrophobic-hydrophilic border for Type I and III rhodopsins in the cell membrane, relative to the position of carotenoid. Prediction was done using PPM 3.0 server ^77^. On the left is the structure of the representative of Type I rhodopsin, XR (PDB ID: 3DDL ^74^). On the right is the structure of the representative of Type III rhodopsin, Cv06 (PDB ID: 9VFC). Retinal marked red; external carotenoids marked orange; protein surfaces colored based on the hydrophility/hydrophobity of amino acid residues.

Thus, the analysis suggests that the “orientation” of CBRs strictly depends on whether the carotenoid is delivered from inside or outside the cell. The exact mechanism of xanthophyll insertion into the membrane is not well understood ^55^. However, it has been suggested that carotenoid-binding proteins deliver carotenoids synthesized in the cytoplasm to the lipid monolayer closest to the site of delivery ^54^. The same could be expected when carotenoids are delivered via mixed micelles. It explains a crucial role of retinal asymmetrical position in the rhodopsins, the second antenna binding to the protein surface.

Indeed, upon insertion into the membrane, carotenoids orient themselves such that their polar groups remain exposed to the aqueous phase, anchoring the molecule at the side of membrane entry. Given that the length of the carotenoid molecule (∼29 Å) closely matches the thickness of the hydrophobic core of the lipid bilayer (∼35 Å), and that the polyene chain typically forms a small angle with the membrane normal ^56,57^, the unanchored keto ring is expected to extend beyond the opposite side of the hydrophobic core and into the CBS. Taken together, these structural features suggest that the CBS is positioned on the membrane side opposite the carotenoid delivery point. This conclusion is further supported by membrane modeling of CBRs positioning (Fig. 6).

Collectively, these observations suggest that HeRs may confer a selective advantage for phototrophy by enabling the use of exogenously acquired secondary antenna systems, thereby expanding the ecological and functional versatility of light-driven energy capture on Earth.

## Discussion

Heliorhodopsins (HeRs) remain enigmatic, particularly regarding their biological roles and molecular functions. While some studies have suggested possible functions, high-resolution structural data exist for only two HeRs, and even these are limited to their ground states. Our study presents the first comprehensive structural and functional characterization of eukaryotic HeR, Cv06, addressing a key gap in our understanding and highlighting structural differences across HeRs from various domains of life.

The central part of our work addresses two major longstanding questions of HeRs biology: (i) their apparent restriction to monoderm organisms, and (ii) their inverted membrane topology relative to Type I rhodopsins. The identification of the fenestration on the surface Cv06 initiated a comprehensive analysis of their presence in other HeRs. The fenestration appeared to be conserved CBS, structurally homologous with that of two-antenna rhodopsins. It was revealed by a complementary study that used docking and bioinformatic analyses. This resulted in a unifying hypothesis: HeRs function as CBRs, utilizing exogenous carotenoids as secondary light-harvesting antennas.

A key structural insight is the presence of a lateral fenestration in Cv06, homologous to CBS in two-antenna Type I rhodopsins such as XR ^21,22^ and Kin4B8 ^25^. Despite differences in membrane orientation and phylogenetic origin, HeRs and Type I CBRs share conserved amino acid residue configurations at the CBS, suggesting functional convergence. Bioinformatic analyses reveal that lateral fenestrations are present in 91.3% of HeRs – substantially more than in Type I rhodopsins. Docking studies further demonstrate that environmental carotenoids, including zeaxanthin, can be positioned on the surface of Cv06 in energetically favorable poses, reinforcing the hypothesis that HeRs can utilize exogenous carotenoids as antenna pigments.

The absence of HeRs in canonical diderm organisms is explained by outer membrane permeability constraints. Diderm outer membranes are impermeable to hydrophobic molecules such as carotenoids, preventing the acquisition of environmental xanthophylls. In contrast, monoderms lack this barrier, enabling the integration of exogenous carotenoids into membrane proteins.

If HeRs function as CBRs requiring exogenous carotenoids, their restriction to monoderm organisms becomes logical. Carotenoids face significant barriers when traversing the outer membrane of diderm cells ^49^, where standard Type I rhodopsins are typically localized ^58,59^. In diderms, the cell wall impedes carotenoid passage, whereas in monoderms, co-localization of HeRs and carotenoids facilitates interaction. Thus, for Type I rhodopsins with CBS, carotenoids must be synthesized internally, necessitating the presence of carotenoid biosynthesis enzymes in these organisms. In contrast, such a requirement is relaxed in monoderm organisms, particularly eukaryotes like algae, which can synthesize and import carotenoids. These findings explain the unique topology and restricted phylogenetic distribution of HeRs and reveal a novel strategy for light harvesting, expanding the known diversity of microbial phototrophy and providing new insights into the evolutionary constraints shaping membrane protein topology.

## Materials and Methods

### Recombinant gene construction and expression cassette preparation

DNA fragment encoding Cv06 (UniProt ID: A0A0G4GC06) was constructed and optimized for expression in the LEXSY expression system (Jena Bioscience) using the GeneArt service (Thermo Fisher Scientific) and was chemically synthesized *de novo*. The fragment flanked by the BglII and KpnI restriction enzyme sites was inserted into the pLEXSY_I-blecherry3 plasmid vector (Jena Bioscience), which encodes a hexahistidine sequence at the C-terminus of the target gene. An expression cassette containing the target gene and nucleotide sequences encoding a zeocin-resistant factor and mCherry co-expressed protein was prepared by hydrolysis of the plasmid vector with the SwaI restriction enzyme.

### Obtaining producer strain

The expression cassette was transfected into LEXSY cells by electroporation. One 0.2 cm gap electroporation cuvette contained 5 μg of the expression cassette and 10^8^ cells/ml resuspended in 500 μl of 5 mM potassium phosphate buffer, pH 7.2, with the addition of 90 mM inositol. Electroporation was performed using MicroPulser Electroporator (Bio-Rad Laboratories) with manual parameters 500 V and monitored pulse time 2.2-2.7 ms. Transfected cells were grown at 26 °C overnight in the dark without shaking in vented tissue culture flasks containing 10 ml of Brain-Heart-Infusion Broth supplemented with 5 μg/ml Hemin, 50 U/ml penicillin, 50 μg/ml streptomycin, 100 μg/ml Nurseotricin, and 100 μg/ml Hygromycin. Then, cells were planted on a BHI agar plate supplemented with 5 μg/ml Hemin, 50 U/ml penicillin, 50 μg/ml streptomycin, 100 μg/ml Nurseotricin, 100 μg/ml Hygromycin, and 100 μg/ml zeocin and incubated at 26 °C in the dark until colonies became visible. After that, the colonies were induced by 10 μg/ml tetracycline and incubated again at 26 °C in the dark until colonies became purple. Colored colonies were checked for the target protein by western blotting.

### Expression of Cv06 in LEXSY membranes

LEXSY culture, producing Cv06, was diluted to OD_600_ = 0.15 by fresh BHI media supplemented with 5 μg/ml Hemin, 50 U/ml penicillin, 50 μg/ml streptomycin, and was shaken at 26 °C in the dark with 140 rpm in non-baffled flasks for about 24 h until OD_600_ reached 1.5 units. Then the culture was induced by 10 μg/ml tetracycline and incubated with the same parameters for 48 h more. After that, cells containing the target protein were collected by centrifugation at 5000 g for 10 min.

### Cv06 extraction and purification

The LEXSY cells expressing Cv06 HeR were resuspended in 20 mM Na-phosphate buffer, pH 7.5, 150 mM NaCl, 1 mM PMSF, and 1 mM EDTA and then were disrupted using a high-pressure homogenizer. The membrane fraction was collected by centrifugation for an hour at 70000 g and 4 °C. Then it was resuspended again in 20 mM Na-phosphate buffer, pH 7.5, 500 mM NaCl, 1 mM PMSF, and 0.5 mM EDTA and centrifuged with the same parameters. Washed membranes were solubilized overnight in 1% (w/v) n-dodecyl β-D-maltoside detergent (DDM), 20 mM Na-phosphate, pH 7.5, 500 mM NaCl, in the presence of 10 μM all-*trans*-retinal. The solubilized protein was centrifuged for an hour at 70000 g and 4 °C, the supernatant was applied to a Ni-affinity resin (Ni-NTA agarose, Qiagen) and eluted with 20 mM Na-phosphate, pH 7.5, 500 mM NaCl, 250 mM imidazole, 10% (v/v) glycerol and 0.05% (w/v) DDM. Then the protein was purified by size exclusion chromatography using Superdex 200 column (GE Healthcare) in 20 mM Na-phosphate, pH 7.5, 500 mM NaCl, 10% (v/v) glycerol, and 0.05% (w/v) DDM. The purified protein was concentrated to 20 mg/ml by centrifugation at 4000 g and 4 °C with the 30kDa Amicon Ultra filter (Millipore).

### pH Titration

The concentrated solution of Cv06 was diluted to 0.5 mg/ml by 10 mM sodium citrate, 10 mM MES, 10 mM HEPES, 10 mM MOPS, 10 mM CHES, 10 mM CAPS, pH 8.0, 300 mM NaCl, 5% (v/v) glycerol, and 0.05% (w/v) DDM. The sample pH was adjusted by concentrated solutions of HCl or NaOH. The absorption spectrum of the sample at each pH was measured using a UV−vis spectrometer (UV-2450, Shimadzu).

### Laser-Flash Photolysis

Cv06 protein solution was suspended in the buffer 20 mM Na-phosphate (pH 7.5), 500 mM NaCl, 10% glycerol, 0.05% (w/v) DDM at a final absorption value at 530 nm of around 0.6 a.u. The sample was excited by a 550 nm nanosecond pulsed Nd;YAG laser (Surelite I-10, Continuum) and an optical parametric oscillator (Surelite OPO Plus, Continuum), as detailed previously. In synchronization with the laser activation, the sample was illuminated with a probe light, monochromated output of a halogen lamp through a monochromator (CT-10, Jasco). For recording time-resolved absorption changes at specific wavelengths, the intensity of the probe light passing through the sample before and after the excitation was monitored by a side-on photomultiplier tube (H8249-102, Hamamatsu Photonics). To improve the signal-to-noise ratio, measurements of the absorption changes were repeated more than 10 times and then averaged. Signals within the first 10 μs after the excitation were ignored due to the presence of the scattered laser light. The sample temperature in a plastic cuvette was kept at 25 °C using a thermostat.

### Proteoliposomes Preparation

Liposomes were produced from asolectin from soybean (Sigma). To prepare 1 ml of proteoliposomes, 10 mg of asolectin was dissolved in 1 ml of chloroform in a pear-shaped glass flask. The flask was rotated using a rotary evaporator at 110 rpm under vacuum until the total evaporation of the solvent and the formation of a thin lipid film on the sides of the flask. Residual chloroform was removed using a vacuum pump for 3 h. Dried asolectin was resuspended by vortexing in 10 mM Tris, 10 mM MES pH 7.2, 150 mM NaCl, supplemented with 2% (w/v) sodium cholate. The protein was reconstituted into liposomes by mixing the liposomes with protein solution and removing detergent using Bio-Beads SM-2 absorbent (Bio-Rad). The lipid mixture was sonicated at 4 °C for 5 min and centrifuged for 5 min in a tabletop centrifuge. The target protein in DDM micelles was added to the supernatant. Then Bio-Beads were immediately added to the mixture and the mixture was rotated for 1 h at 4 °C. Then the used beads were replaced with new ones and the procedure was repeated 5 times.

### Planar bilayer lipid membrane experiments (BLM)

The planar bilayer lipid membrane was formed with a solution of 1,2-diphytanoyl-sn-glycero-3-phosphocholine and 1,2-dimyristoyl-sn-glycero-3-ethylphosphocholine in n-decane (20 and 0.4 mg/ml) on a round aperture with a diameter of 0.6 mm, separating two identical sections of the Teflon cell. Each section was filled with 1.5 ml of buffer 10 mM Tris, 10 mM MES pH 7.2, 150 mM NaCl, and the aperture was below the liquid level. The electrical current was measured with two AgCl electrodes placed into the solutions on the two sides of the BLM via agarose bridges, using a DLPCA-200 amplifier (FEMTO® Messtechnik GmbH, Germany). 30 µl of proteoliposomes were added to one section and incubated for 1 hour with constant stirring. Then, 0.5 µM of protonophore 4,5,6,7-tetrachloro-2-trifluoromethylbenzimidazole (TTFB) was added to another section. In one hour, BLM was illuminated with a halogen lamp (Olympus KL2500-LCD, 250 W).

### Molecular docking assay

Molecular docking was performed using Gnina v.1.0 software with exhaustiveness = 32 ^60,61^. Ligand molecules for docking were prepared with DataWarrior v.06.01.00 software using option “Generate conformers” with the following input parameters: algorithm – random; low energy bias; forcefield – MMFF94s+; maximum conformer count – 1 per stereoisomer ^62^. Ligand molecules were then processed to produce PDBQT format using OpenBabel v.3.1.1 ^63^.

### Crystallization, data collection, and structure refinement

The crystals of Cv06 were grown with an *in meso* approach ^64^, similar to that used in our previous works ^36–38,40,43^. Briefly, the solubilized protein was concentrated to 40 mg/ml and was mixed with the premelted at 42 °C monopalmitolein (Nu-Chek Prep) in a 3:2 ratio (lipid:protein) to form a lipidic mesophase. The mesophase was homogenized in coupled syringes (Hamilton) by transferring the mesophase from one syringe to another until a homogeneous and gel-like material was formed. 100 nl drops of a protein–mesophase mixture were spotted on a 96-well LCP glass sandwich plate (Marienfeld) and overlaid with 500 nL of a precipitant solution using the NT8 crystallization robot (Formulatrix). The crystals were grown at 20 °C and appeared in 4 months.

The best diffracting crystals were obtained in 1.2 M sodium potassium phosphate buffer, pH 8.2. The crystals were cryoprotected with 10% (v/v) glycerol and 2.16 M sodium potassium phosphate buffer (pH 8.2), harvested using MicroMounts (MiTeGen), flash-cooled, and stored in liquid nitrogen before data collection.

X-ray diffraction data were collected at the ID23-2 beamline of the European Synchrotron Radiation Facility, Grenoble, France, at 100 K. Data collection protocol was optimized with BEST ^65^. Diffraction images were integrated with XDS software ^66^ and anisotropically scaled with STARANISO web service ^67^ (https://staraniso.globalphasing.org/cgi-bin/staraniso.cgi). The criterion for the anisotropic cutoff was local I/σI > 0.5. The diffraction limits were 2.5×2.4×3.0 Å. The data collection statistics are shown in Table S1.

MoRDa automatic molecular replacement pipeline ^68^ from CCP4 online web service ^69^ was used to solve the phase problem. The crystal structure of *T*aHeR (PDB 6IS6 ^13^) was chosen as a reference ^13^. Molecular replacement was successful in the P1 space group and contained four dimers of Cv06 in the asymmetric unit. The model was iteratively refined using Phenix.Refine ^70^, and Coot ^71^. The quality of the final model was assessed using Phenix.Molprobity ^72^. Paired refinement technique ^73^ determined the final resolution of the model, which dropped to 2.6×2.6×3.0 Å. The refinement statistics are shown in Table S1.

## Acknowledgments

The diffraction experiments were performed at the ID30B, ID30A-3, ID23-1, and ID23-2 beamlines of the European Synchrotron Radiation Facility (ESRF), Grenoble, France (proposals MX-2270 and MX-2330). We are grateful to local contacts at the ESRF, especially Alexander Popov and Igor Melnikov, for aiding in using these beamlines. V.G. was supported by grant ANR-19-CE11-0026. DK, FT, EZ, VB, and PK acknowledge the Ministry of Science and Higher Education of the Russian Federation (agreement # 075-03-2025-662, project FSMG-2024-0012). Crystallography data treatment was supported by the Ministry of Science and Higher Education of the Russian Federation (agreement # 075-15-2025-512). The characterization of protein spectral kinetics and photocycles was supported by the Russian Science Foundation (RSF), project 21-64-00018.

## Author Contributions

V.Gor. conceived the project. D.K. and V.Gol. performed expression, purification, and pH titration. A.M. and S.Bukha performed BLM. F.T. carried out laser-flash photolysis. S.Bukhd., R.A., and M.R. performed crystallization. S.Bukhd. and M.S. solved and refined the structure. D.K. and S.I. conducted molecular docking. D.K. and E.Z. carried out bioinformatic analyses. D.K., S.Bukhd., and V.Gor. wrote the original draft. V.Gor., E.B., and V.B. supervised the project. P.K., Y.R., and A.V. managed project administration. All authors discussed the results and reviewed the final version of the manuscript.

## Data Availability

X-ray crystallography atomic coordinates and structure factors for Cv06 heliorhodopsin have been deposited in the Protein Data Bank under accession code 9VFC. Data supporting the findings of this manuscript are available from a corresponding author upon reasonable request.

