## Supplementary Information for "Topological adaptation of heliorhodopsins enables exogenous second-antenna acquisition in monoderm phototrophs"

**This PDF file includes:**

Figures S1 to S8
Table S1
SI References

### Figures

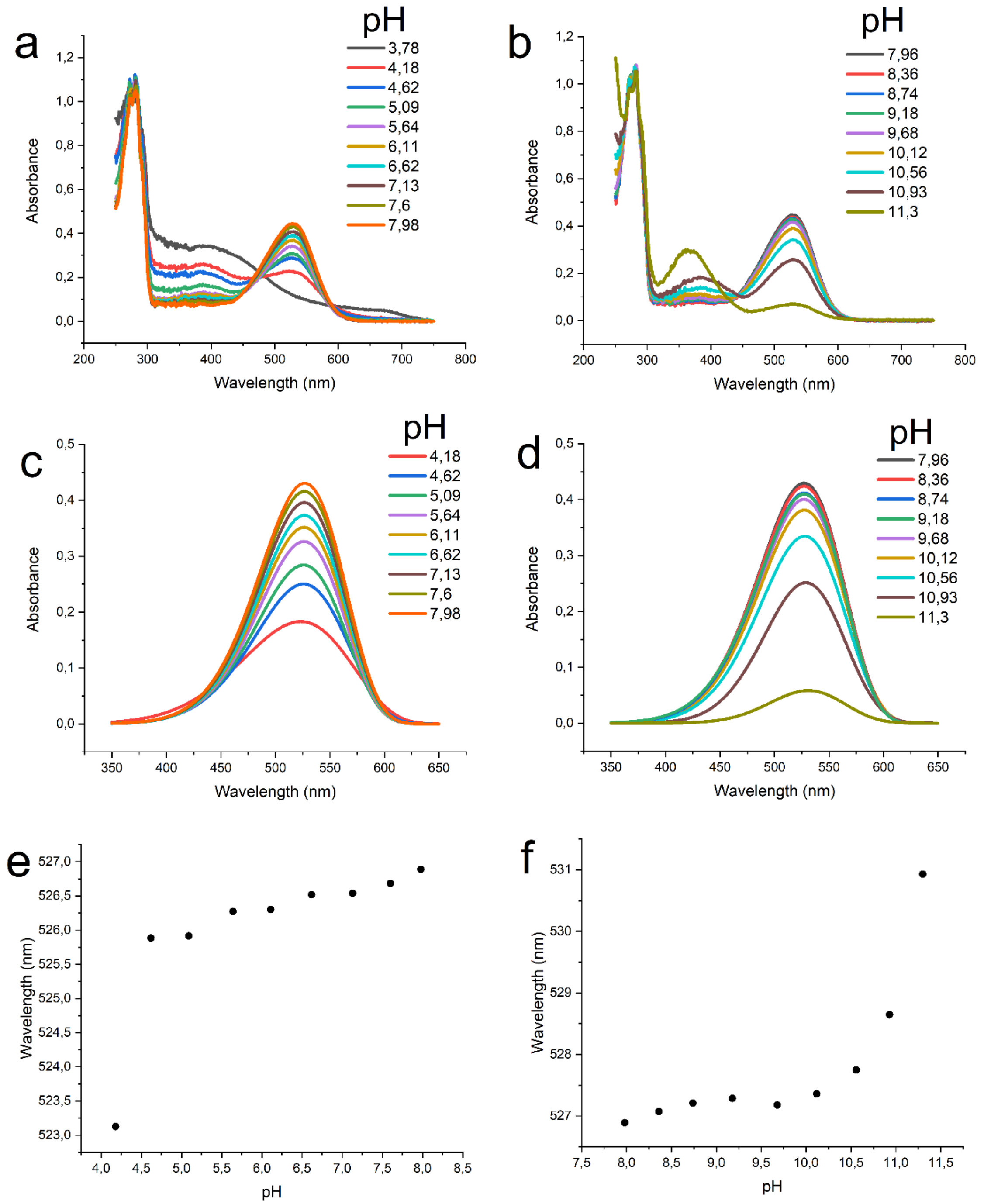


**Fig. S1. Cv06 pH titration experiments.** Absorption spectra of Cv06 in acidic (**a**) and alkaline (**b**) environments (raw data). Fitted rhodopsin peaks of Cv06 in acidic (**c**) and alkaline (**d**) environments. Absorption extremum relation of rhodopsin peak and pH of solution: acidic (**e**) and alkaline (**f**) areas. Measurements were performed using starting buffer 10 mM sodium citrate, 10 mM MES, 10 mM HEPES, 10 mM MOPS, 10 mM CHES, 10 mM CAPS, pH 8.0, 300 mM NaCl, 5% (v/v) glycerol, and 0.05% (w/v) DDM.

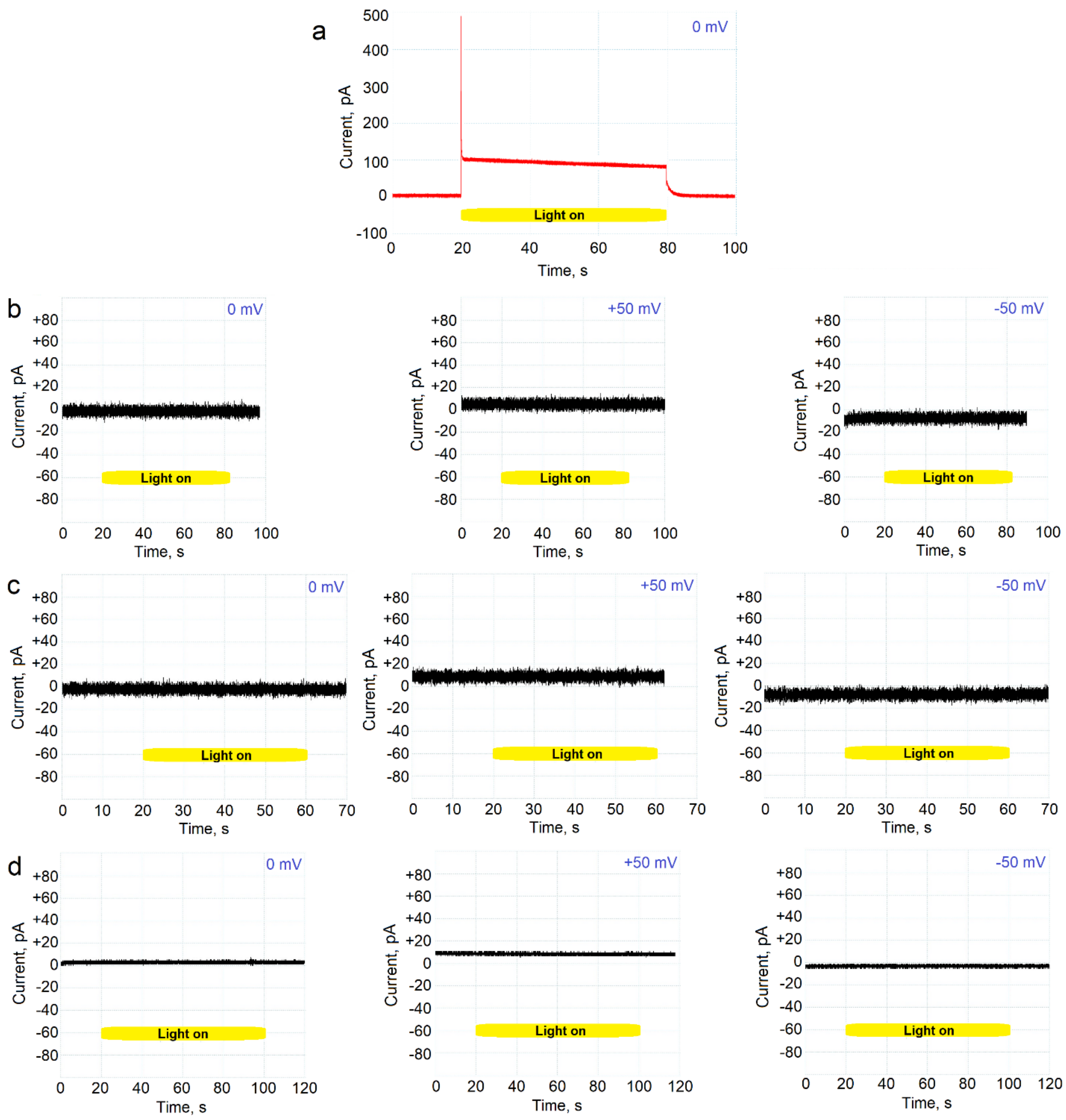


**Fig. S2. BLM essay of Cv06 in lipid vesicles.** Ion current on lipid membrane with integrated vesicles containing examined rhodopsin during the time depending on illumination. (**a**) BR, as a positive control, is marked red. (**b**) Cv06 in the buffer 10 mM Tris, 10 mM MES pH 7.2, 150 mM NaCl containing $H^+$, $Na^+$, $K^+$, $Cl^-$ and $H_3COO^-$ ions. (c) Cv06 in buffer 10 mM Tris, 10 mM MES pH 7.2, 150 mM NaCl with the addition of 1 mM $NaNO_3$. (d) Cv06 in buffer 10 mM Tris, 10 mM MES pH 7.2, 150 mM NaCl with the addition of 1 mM $NaHCO_3$. Measurements were conducted for a set of pHs, but are shown only for pH 7.2. For other pHs results are the same as for pH 7.2.

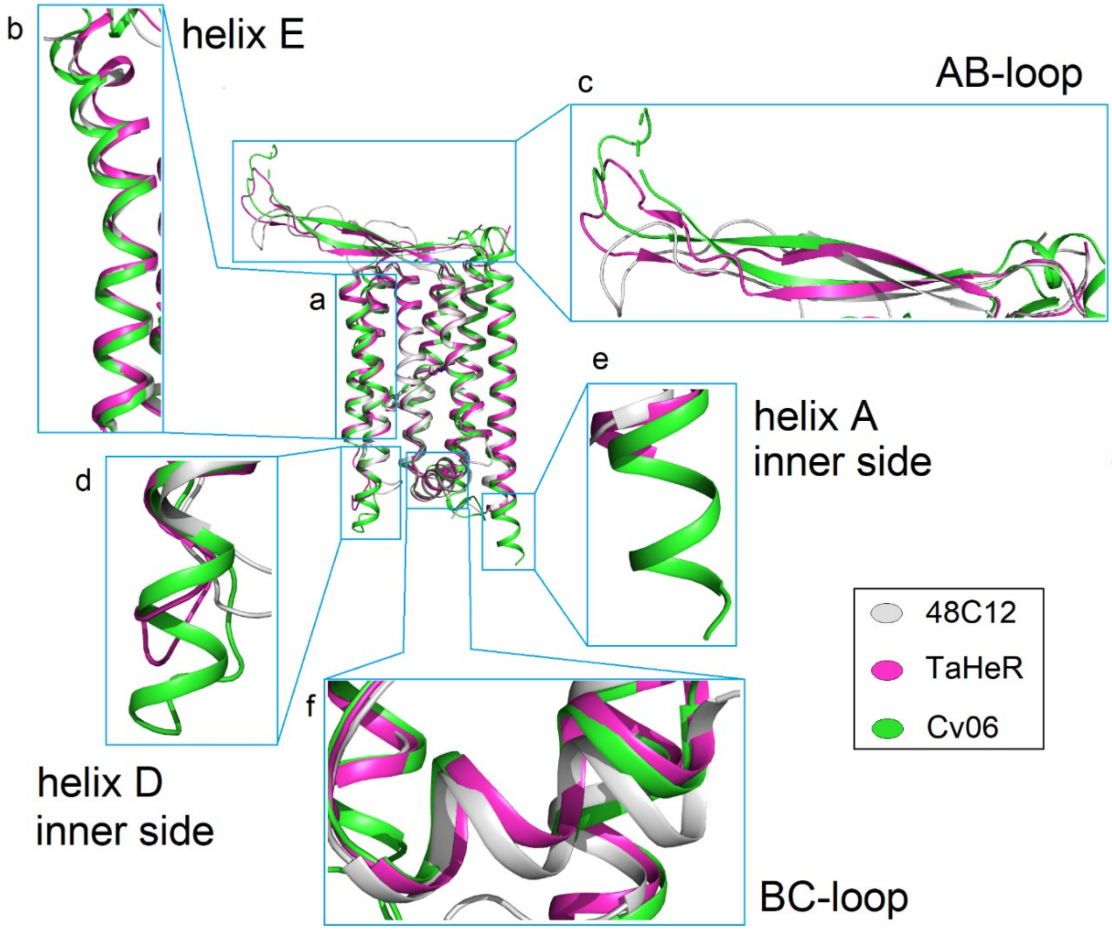


**Fig. S3. Overview of available HeR structures.** (**a**) Superposition of 48C12 (PDB ID: 6SU3 [1], chain X), *Ta*HeR (PDB ID: 6IS6 [2], chain A) and Cv06 (PDB ID: 9VFC, chain A); (**b**) helices E; (**c**) AB-loops; (**d**) inner side of helices D; (**e**) inner side of helices A; (**f**) BC-loops. (**a-f**) 48C12 marked grey, *Ta*HeR marked pink, and Cv06 marked green.

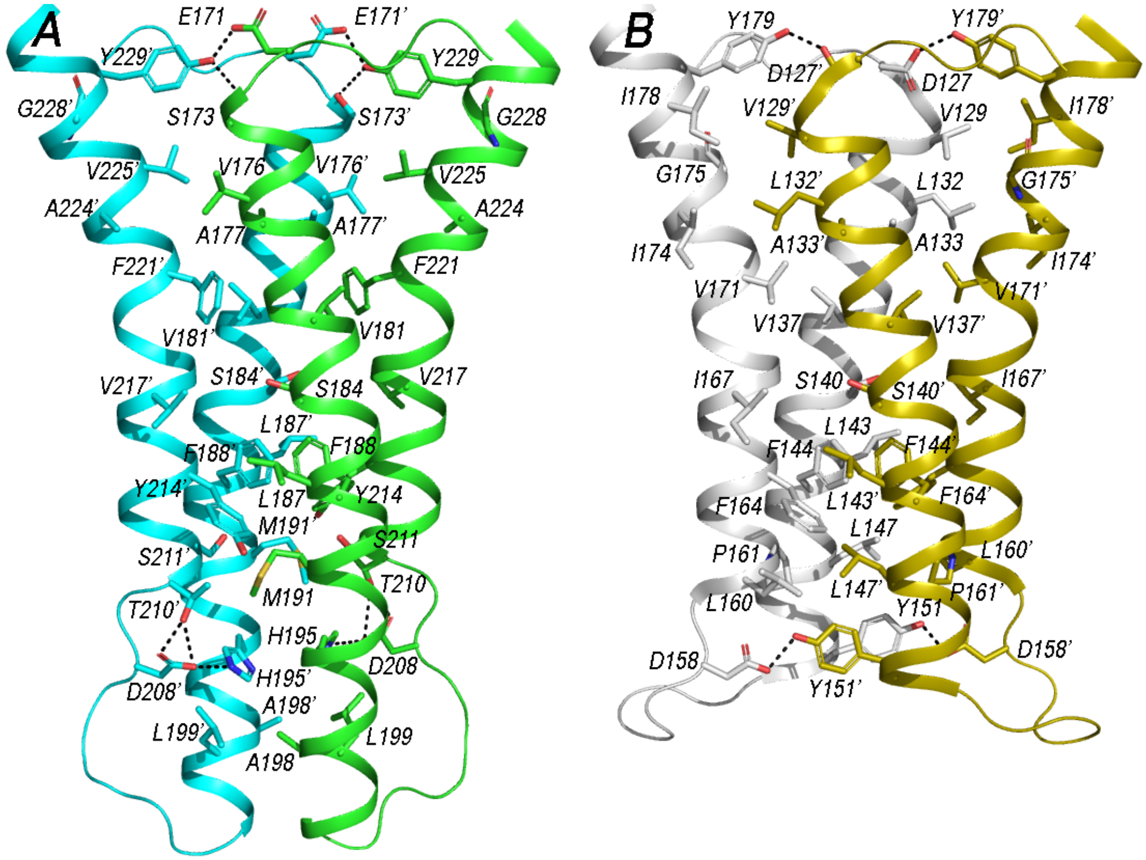


**Fig. S4. Comparison of the dimerization interface in Cv06 and 48C12.** Separate protomers of (**A**) Cv06 (PDB ID: 9VFC) and (**B**) 48C12 (PDB ID: 6SU3 [1]) are shown in different colors.

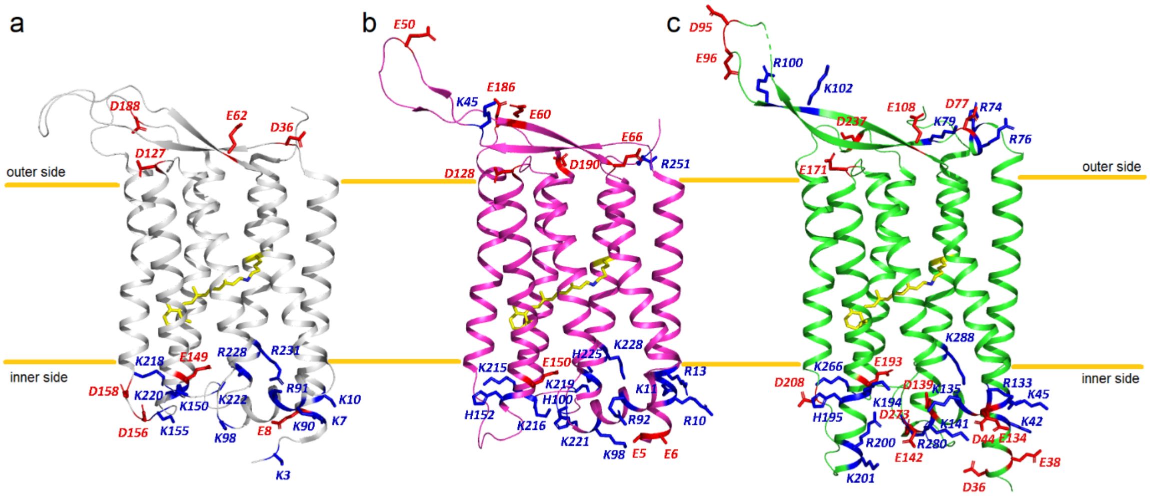


**Fig. S5. Topology of HeRs.** (**a**) 48C12 (PDB ID: 6SU3 [1]) marked grey, (**b**) *Ta*HeR (PDB ID: 6IS6 [2]) marked pink, and (**c**) Cv06 (PDB ID: 9VFC) marked green. Charged amino acid residues are marked blue (positive) and red (negative). Retinal, bound to lysine residue, marked yellow.

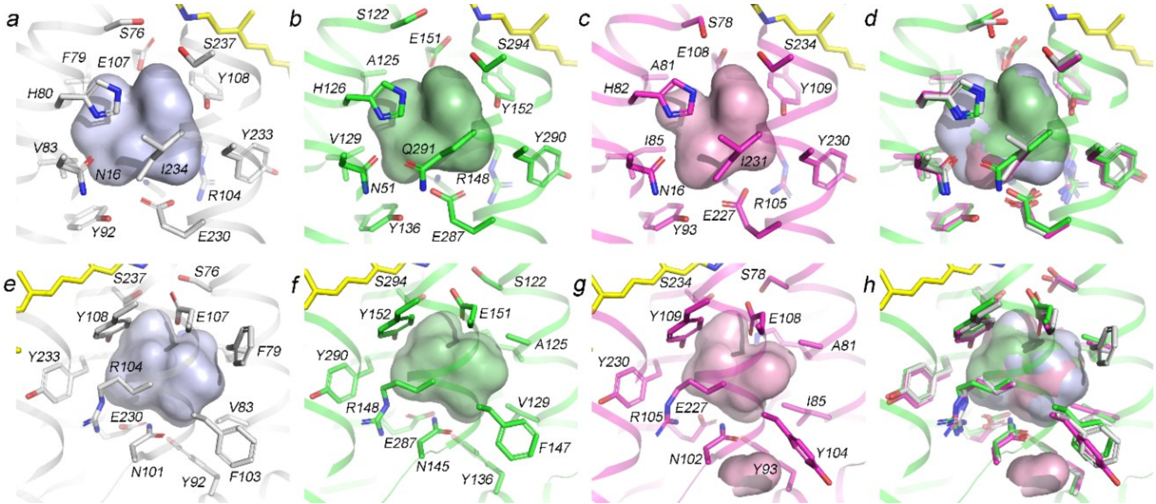


**Fig. S6. Overview of amino acid residues forming large hydrophilic cavities in HeRs.** 48C12 (PDB ID: 6SU3 [1]) marked grey, *Ta*HeR (PDB ID: 6IS6 [2]) marked pink and Cv06 (PDB ID: 9VFC) marked green. Retinal cofactor marked yellow. (**a-c**, **e-g**) Separate cavity demonstrations of 48C12 (**a** and **e**), Cv06 (**b** and **f**), and *Ta*HeR (**c** and **g**). (**d** and **h**) Imposition of the HeRs cavities. The retinal molecule and cartoon backbone are shown only for Cv06. Amino acid residues marked green for Cv06, gray for 48C12, and magenta for *Ta*HeR.

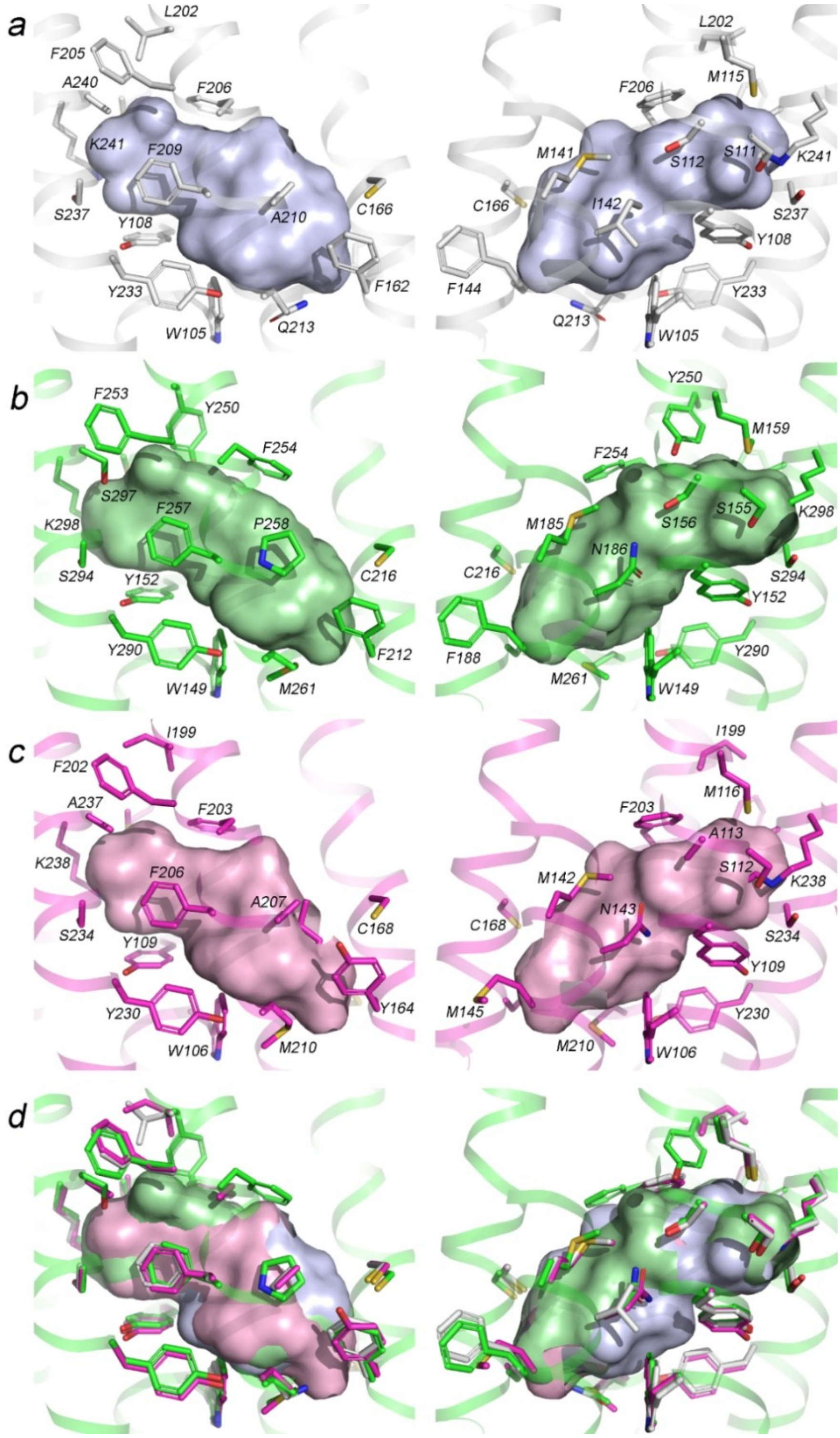


**Fig. S7. Overview of amino acid residues forming the retinal pocket in HeRs.** 48C12 (PDB ID: 6SU3 [1]) marked grey, *T*aHeR (PDB ID: 6IS6 [2]) marked magenta, and Cv06 (PDB ID: 9VFC) marked green. (**a-c**) Separate retinal pocket demonstrations of 48C12 (**a**), Cv06 (**b**), and *T*aHeR (**c**). (**d**) Imposition of the retinal pockets of bacterial (48C12, gray), archaeal (*T*aHeR, magenta), and eukaryotic (Cv06, green) HeRs. The backbone is shown only for Cv06.

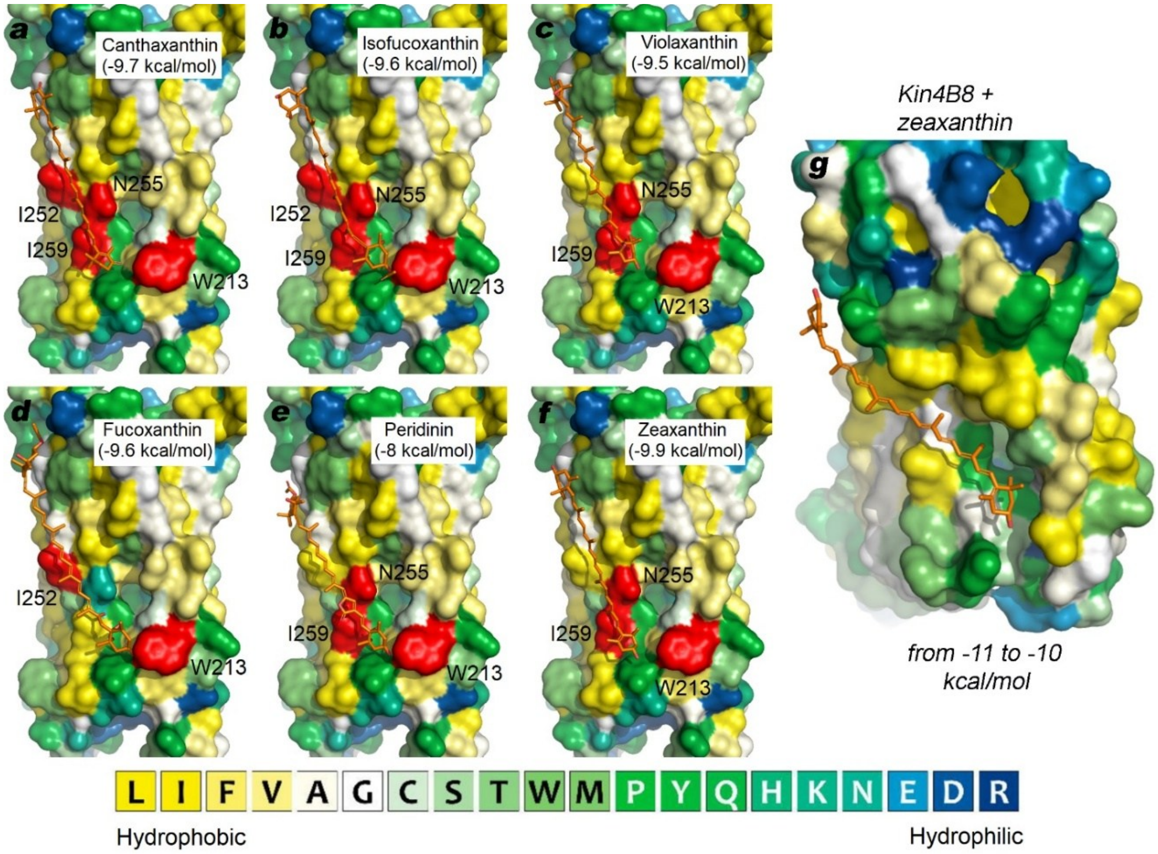


**Fig. S8. Molecular docking assay of Cv06 and carotenoid molecules.** Cv06 surface marked according to the hydrophobicity of corresponding residues. Carotenoids are marked orange. Residues allowed to be flexible during the docking assay are marked red and labeled. The figure shows the most relevant positions of carotenoids: (**a**) canthaxanthin, (**b**) isofucoxanthin, (**c**) violaxanthin, (**d**) fucoxanthin, (**e**) peridinin, (**f**) zeaxanthin. Panel (**g**) shows the experimentally determined position of zeaxanthin on the surface of Kin4B8 (PDB ID: 8I2Z [3]) for comparison.

**Tables**

**Table S1. Data collection and refinement statistics**

| | Cv06 (PDB ID: 9VFC) |
|---|---|
| **Data collection** | |
| Beamline | ESRF ID23-2 |
| Wavelength (Å) | 0.8731 |
| Space group | P 1 |
| Cell dimensions | |
| *a*, *b*, *c* (Å) | 77.89, 77.89, 135.99 |
| a, b, g (°) | 105.67, 101.16, 90.02 |
| Resolution (Å) | 42.98 – 2.50 (2.63 – 2.50) * |
| $R_{pim}$ | 0.121 (1.144) |
| $CC_{1/2}$ | 0.990 (0.168) |
| *I* / s*I* | 4.9 (0.7) |
| Completeness spherical (%) | 78.7 (27.0) |
| Completeness ellipsoidal (%) | 91.8 (69.0) |
| Redundancy | 2.9 (3.2) |
| **Refinement** | |
| Resolution (Å) | 42.98 – 2.60 |
| No. reflections work / free | 79126/1914 |
| $R_{work}$ / $R_{free}$ | 0.2304/0.2713 |
| No. atoms | |
| Protein | 16825 |
| Retinal | 160 |
| Lipids | 1406 |
| Solvent | 153 |
| *B*-factors (Å$^2$) | |
| Protein | 62.51 |
| Retinal | 48.44 |
| Lipids | 59.82 |
| Solvent | 55.10 |
| Ramachandran plot (%) | |
| Favorable | 97.69 |
| Outliers | 0 |
| R.m.s. deviations | |
| Bond lengths (Å) | 0.002 |
| Bond angles (°) | 0.42 |

*Values in parentheses are for highest-resolution shell.
